# Engineering, production and characterization of Spike and Nucleocapsid structural proteins of SARS–CoV-2 in *Nicotiana benthamiana* as vaccine candidates against COVID-19

**DOI:** 10.1101/2020.12.29.424779

**Authors:** Tarlan Mamedov, Damla Yuksel, Merve Ilgın, İrem Gürbüzaslan, Burcu Gulec, Gulshan Mammadova, Deniz Say, Gulnara Hasanova

## Abstract

The COVID-19 pandemic, which is caused by SARS-CoV-2 has rapidly spread to more than 216 countries and has put global public health at high risk. The world urgently needs a cost-effective and safe SARS-CoV-2 coronavirus vaccine, antiviral and therapeutic drugs to control the COVID-19 pandemic. In this study, we engineered the Nucleocapsid (N) and Spike protein (S) variants (Receptor binding domain, RBD and S1 domain) of SARS-CoV-2 genes and produced in *Nicotiana benthamiana* plant. The purification yields were at least 20 mg of pure protein/kg of plant biomass for each target protein. The S protein variants of SARS-CoV-2 showed specific binding to angiotensin converting enzyme 2 (ACE2), the SARS-CoV-2 receptor. The purified plant produced N and S variants were recognized by N and S protein specific monoclonal and polyclonal antibodies demonstrating specific reactivity of mAb to plant produced N and S protein variants. In addition, IgG responses of plant produced N and S antigens elicited significantly high titers of antibody in mice. This is the first report demonstrating production of functional active S1 domain and Nucleocapsid protein of SARC-CoV-2 in plants. In addition, in this study, for the first time, we report the co-expression of RBD with N protein to produce a cocktail antigen of SARS-CoV-2, which elicited high-titer antibodies compared to RBD or N proteins. Thus, obtained data support that plant produced N and S antigens, developed in this study, are promising vaccine candidates against COVID-19.

## Introduction

The novel coronavirus, currently designated as SARS-CoV-2, is a novel and highly pathogenic coronavirus, and spread more than to 222 countries and territories in a short time, and as of December 25, 2020, more than 79,931,215 cases were recorded, and more than 1,765,265 confirmed deaths. Although a number of various types vaccines (mRNA, DNA, and viral vector-based, protein based subunits, inactivated, attenuated etc.) for COVID-19 is underway and there are currently no drugs, no specific treatment and no vaccines available to treat COVID-19 infections and protect people against fatal SARS-CoV-2 coronavirus. The world urgently needs a safe and effective SARS-CoV-2 coronavirus vaccine, antiviral and therapeutic drugs, cost effective diagnostic reagents and kits to control the COVID-19 pandemic and relieve the human suffering associated with the pandemic that kills thousands of people every day. One of the major observed features of this virus is that SARS-CoV-2 is transmitted from infected people without symptoms, therefore, it increases the challenges of controlling a deadly pandemic without the use of a vaccine. One-third of the virus genome (~30 kb) of SARS-CoV encode mainly structural proteins such as spike (S) glycoprotein, nucleocapsid protein (N), small envelop protein (E) and matrix protein (M). When reported on January 2020, in Spike protein, SARS-CoV-2 had about 79.6 % sequence identity with that of SARS-CoV^1,2^. Now the sequence identity is less than 76%, as the spike protein of the new coronavirus SARS-CoV-2 has acquired more than 725 mutations. S protein of coronavirus consist of S1 and S2 domains and plays a key role in virus binding, fusion and entry to host cells. S protein has been demonstrated as a leading target for vaccine and neutralizing antibody development. In fact, most of vaccines, such as mRNA, DNA, viral vector-based, subunits, protein based vaccines against COVID-19 have been developed based on the gene encoding S protein^3–7^. S1 domain contains a receptor-bindingdomain (RBD), which binds specifically to angiotensin-converting enzyme (AEC2), areceptorfor both SARS-CoV and SARS-CoV-2.The S protein of SARS-CoV-2 coronavirus is cycteine-rich protein and nine cysteine residues are found in the RBD, eight of which involve in forming four pairs of disulfide bridges^8^. In addition, the S protein has 22 potential N-glycosylation sites, of which two of them are in the RBD^9^. Thus, correct formation of disulfide bridges and status of glycosylation would be essential for functional activity of recombinant S protein-based vaccines, produced using different expression system. The structure of SARS-CoV-2 spike RBD was recently determined by crystal structure analysis at 2.45 Å resolution^1^. It was demonstrated that the overall ACE2-binding mode of the SARS-CoV-2 RBD is nearly identical to that of the SARS-CoV RBD, which also utilizes ACE2 as the cell receptor. It should be noted that since the RBD is the critical region for receptor binding, therefore RBD based vaccines could be great promise for developing vaccines and also highly potent cross-reactive therapeutic agents. It has been recently demonstrated that RBD-Fc-based COVID-19 vaccine candidate elicited high titer of RBD-specific antibodies with neutralizing activity against SARS-CoV-2 infections^3^. As mentioned above, N protein has been shown to be more conserved compared with S protein and share 90% amino acid homology with SARC-CoV and stable with fewer observed mutations^10–13^. Since the N-protein was found to be more conservative and showed strong immunogenicity, the N-protein could be a strong potential candidate for a COVID-19 vaccine^14^.

Numerous studies in recent years have demonstrated plant expression systems as promising expression platforms for the cost-effective, rapid and safe production of various recombinant proteins. Plant expression systems have several advantages over other expression systems currently in use. This system has been successfully used for rapid and cost-effective production of a variety of recombinant proteins, vaccine candidates, therapeutic proteins, enzymes, antibodies etc.^15–22^. Recently, a number of flexible approaches have been developed, which enabled the successfully production of pharmaceutically important complex proteins including human proteins and enzymes in plants^15,19–21^. Thus, plant expression system could be ideal platform for cost effective, safe and rapid production of structural proteins of SARS-CoV-2 as a vaccine candidates against COVID-19. In previous efforts, a number of studies have been conducted on the transgenic (stable) expression of the spike protein of SARS-CoV in plants. N-terminal fragment of SARS-CoV S protein (S1) was produced in tomato and low-nicotine tobacco plants and demonstrated immunogenicity in mice after parenteral or oral administration^23^. Several groups have reported development of S protein recombinant plant-based vaccines against different coronoviruses for oral delivery that elicit protective immunity against virus challenge^24–26^. Recombinant SARS-CoV spike protein was also expressed in plant cytosol and chloroplasts^27^. However, transgenic plant approach has some concerns, which are mainly associated with the long development time. Additionally low target accumulation levels and the possibility of gene flow from transgenic plants to wild types are of concern^28^. Plant transient expression system has a number of advantages over stable expression. Using a transient plant expression system, the production of glycosylated form of RBD SARS-CoV-2 in *Nicotiana benthamiana* plant has been reported recently^29^. However, the expression level of RBD in *N. benthamina* plant was very low, 8 μg/g, which is unsatisfactory to be economical for commercialization. In this study, we report the high production of functionally active structural proteins of SARC-CoV-2 such as RBD (glycosylated and deglycosylated variants), and glycosylated variant of S1 domain, and N proteins in the *N. benthamiana* plant using a transient expression platform. Notably, neither the S1 domain of SARS-CoV nor the SARS-CoV-2 was produced using a transient plant expression system elsewhere.

## Results

### Engineering, cloning and expression, purification and characterization of N andS protein variants in *N. benthamiana* plant

The purification yields were at least 20 mg pure protein/kg plant biomass for each target protein. On SDS-PAGE and western blotting, plant produced N protein molecule appears as a doublet protein with a MM of 48 and 24 kDa (Figure 1). Both molecular forms of plant produced N proteins were recognized by anti-FLAG antibody (Figure 1B) and by N protein specific mAb (Figure 1C), demonstrating specific reactivity of mAb to the two forms and also confirm that epitope for mAb still present in ~24 kDa fragment. N protein of SARS-CoV-2 has 5 potential N-glycosylation sites. When N protein was co-expressed with Endo H, there was no protein band shift in the western blotting suggesting that plant produced N protein is not N-glycosylated. To understand the role of glycosylation, we produced both glycosylated and non-glycosylated variants of RBD and S1 protein in *N. benthamiana* plant. Figure 2 demonstrate SDS-PAGE analysis of the anti-Flag column purified, deglycosylated and glycosylated variants of RBD. The purity of both glycosylated anddeglycosylated proteins were higher than 90%. Plant produced, purified glycosylated RBD molecules appear as a protein with a molecular mass (MM) of ~36 kDa (Figure 2, dRBD). dRBD molecules appear as a double protein with a MM of ~32 or ~33 kDa (Figure 2, dRBD). Plant produced dRBD and gRBD proteins were very well recognized by anti-FLAG (Figure 2A), and also by S protein specific polyclonal (Figure 2C) antibody, demonstrating specific reactivity of pAb to the RBD protein. N+gRBD co-expressed (cocktail) proteins were purified using anti-Flag column. SDS-PAGE analysis of the cocktail protein purified from *N. benthamiana* plant is presented in Figure 3.Deglycosylated variants were produced using the *in vivo* Endo H deglycosylation strategy, which we recently developed^19^. Deglycosylated and glycosylated variants of plant produced S1 proteins were purified using HisPur™ Ni-NTA resin. On SDS-PAGE and western blotting, plant produced glycosylated S1 (gS1) protein appears as a single protein with a MM of ~100 kDa (Figure 4A). No clear visible band of Ni-NTA resin purified, Endo H deglycosylated S1 protein (dS1) was observed in SDS-PAGE gels, suggesting degradation of deglycosylated S1 protein. However, on Western blotting, deglycosylated S1 protein with a band of ~ 80 kDa was observed. Plant produced, purified glycosylated and deglycoslated S1 variants were recognized by anti-His mAb tag (Figure 4B) or anti-S protein specific pAb (Figure 4C).

**Figure 1.**
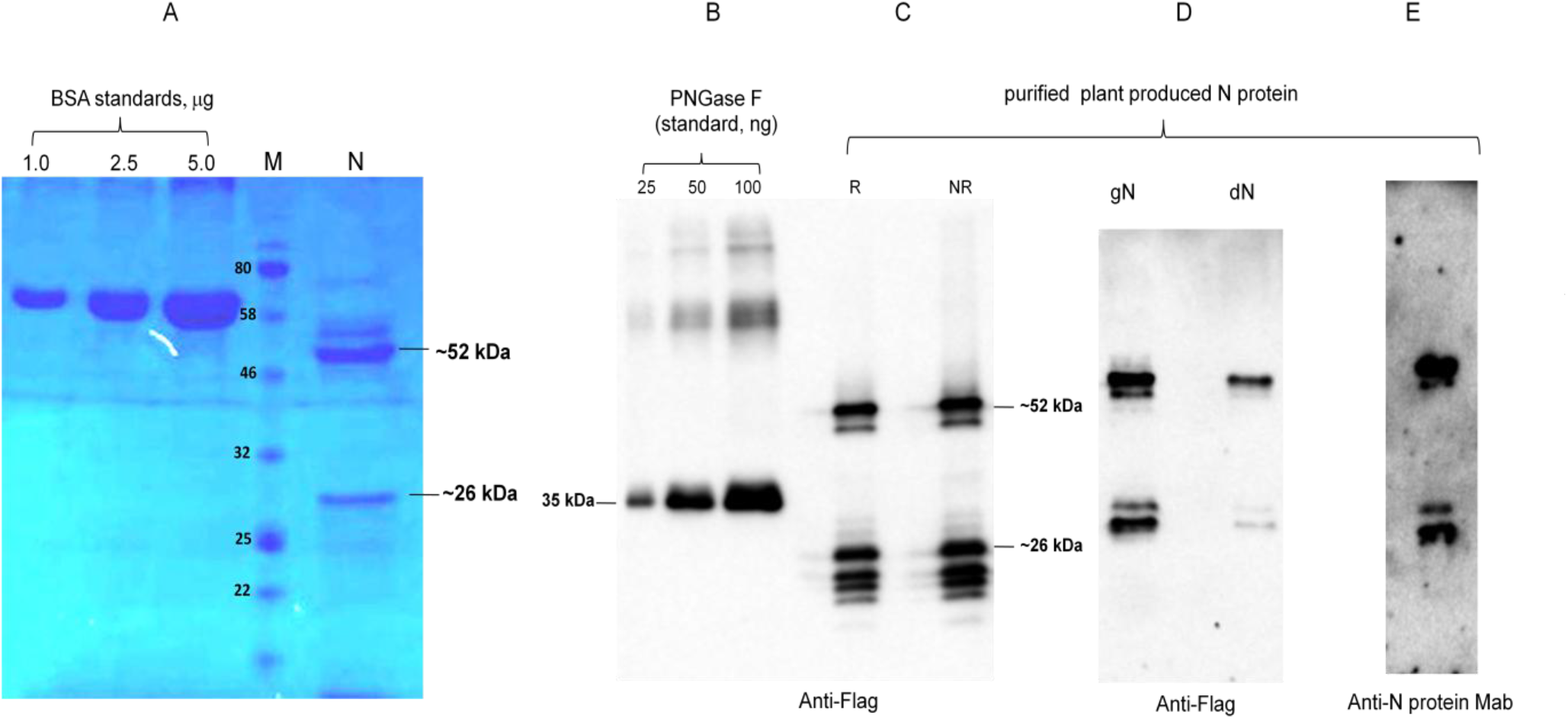
SDS-PAGE (A) and Western blot (B, C) analysis of purified plant produced N protein. N protein from *N. benthamiana* plant was purified using anti-FLAG antibody resin. Membranes were probed with anti-Flag (B) or anti-N protein monoclonal antibody (C). N: 2 μg plant produced; purified N protein was loaded into well. BSA standards: 1.0, 2.5 and 1.5 μg BSA protein was loaded as a standard protein. R: reducing. NR: non-reducing gel. M: color prestained protein standard.

**Figure 2.**
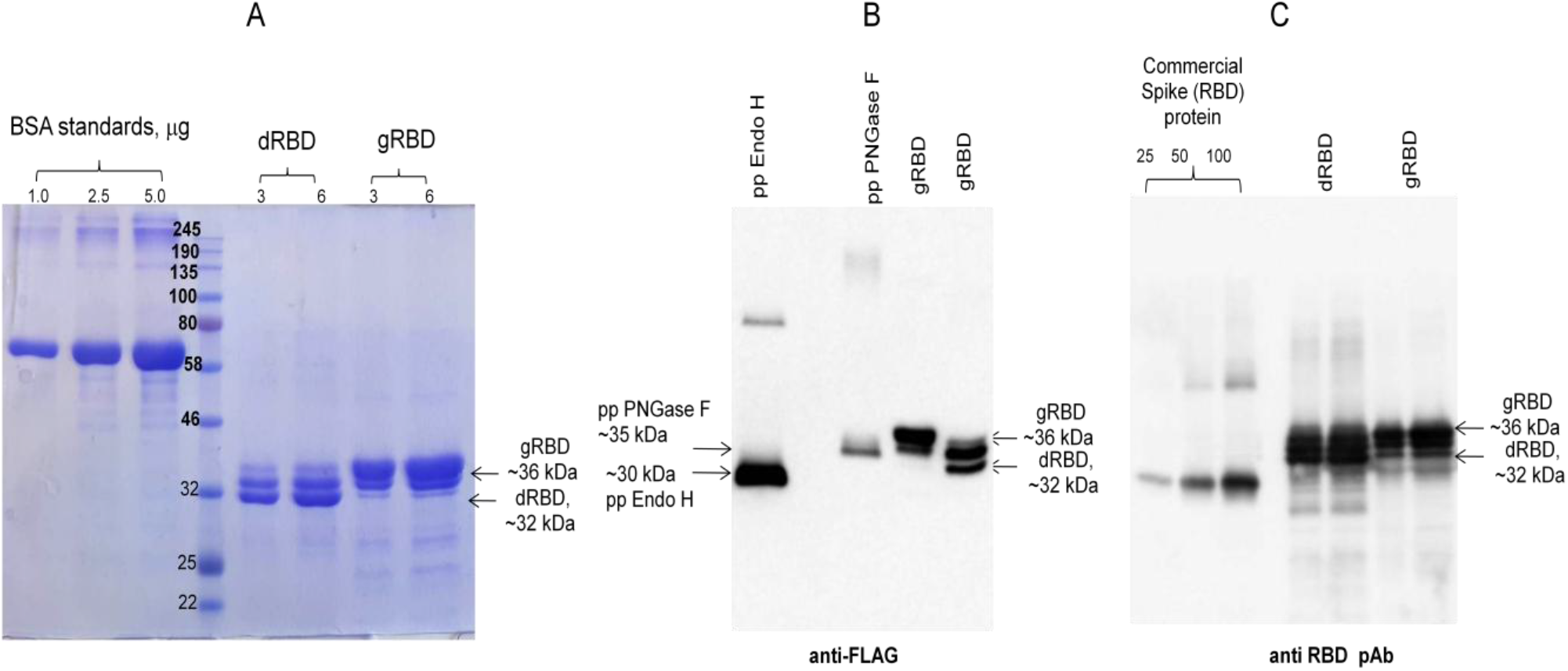
SDS-PAGE analysis of purified plant produced RBD protein (Glycosylated and non-glycoslated) RBD protein variants from *N. benthamiana* plant were purified using anti-FLAG antibody resin. A: gRBD: 3 μg or 6 μg plant produced glycosylated RBD. dRBD: 3 μg or 6 μg plant produceddeglycosylated RBD. BSA standards: 1.0, 2.5 and 5.0μg BSAstandards.M: color prestained protein standard (NEB).B: Western blot analysis of gRBD and dRBD using anti-Flag antibody. pp Endo H, plant produced and purified Endo H protein with Molecular mass of ~30 kDa^19^. pp PNGase F, plant produced and purified PNGase F protein with Molecular mass of ~35 kDa^19^. C: Western blot analysis of gRBD and dRBD using commercial available anti-RBD antibody (MBS2563840, MyBiosource, USA); Commercial spike protein: COVID 19 Spike Protein (RBD) Active Protein; sequence positions Arg319-Phe541, MBS2563882, MyBiosource, Inc, USA).

**Figure 3.**
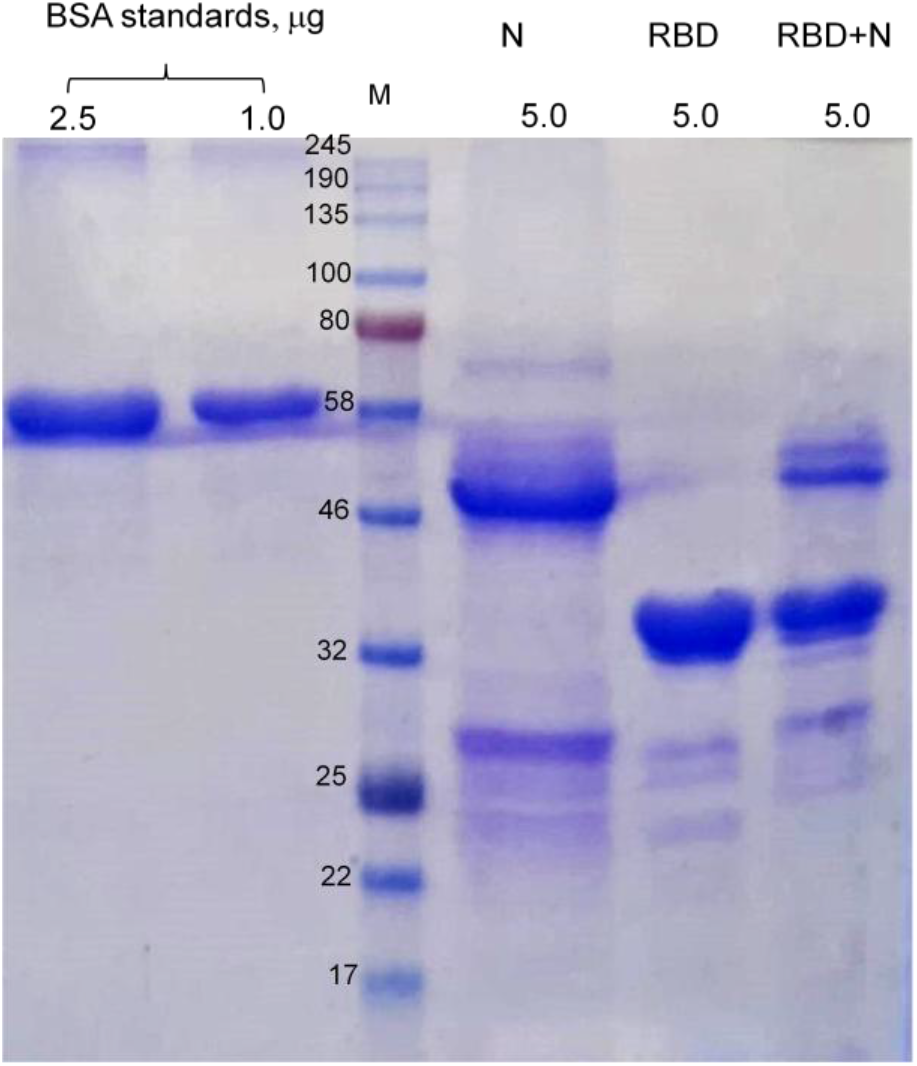
SDS-PAGE analysis of co-expression of RBD with N protein. SDS-PAGE analysis of plant produced, purified N +RND proteins. N: plant produced purified N protein; RBD: plant produced purified RBD protein; RBD+N: plant produced purified RBD+N from co-expression of BRD and N protein. BSA standards: 1.0 and 2.5μg BSA protein was loaded as a standard protein. M: color prestained protein standard (NEB, USA). **Figure 3. Western analysis of purified plant produced S protein (glycosylated)** S protein from *N. benthamiana* plant was purified using anti-FLAG antibody resin. Membranes were probed with anti-FLAG (A), anti-S (RBD) mAb (B) or anti-S (RBD) pAb (C).

**Figure 4.**
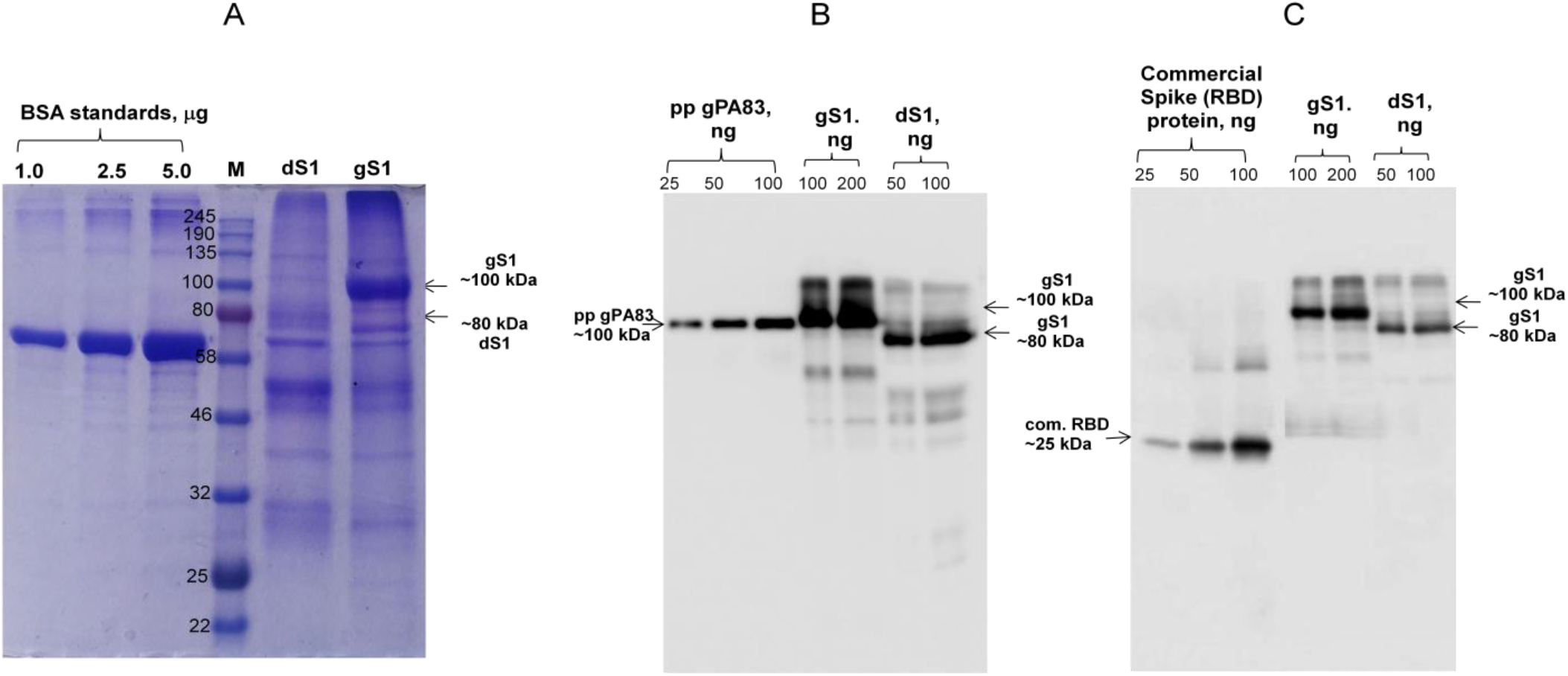
SDS-PAGE (A) and Western blot (B, C) analysis of purified plant produced S1 protein. S1protein variant from *N. benthamiana* plant were purified using HisPur™ Ni-NTA Resin. Membranes were probed with anti-His tag monoclonal antibody (B) or commercial available anti-RBD antibody (MBS2563840, MyBiosource, USA); M: color prestained protein standard (NEB). gS1: plant produced glycosylated S1; dS1: plant produced deglycosylated S1. Commercial spike protein: COVID 19 Spike Protein (RBD) Active Protein; sequence positionsArg319-Phe541, cat. no. MBS2563882, MyBiosource, USA).

### Binding of plant produced RBD and S1 proteins to ACE2

Binding of plant-produced gRBD, dRBD and gS1 proteins to ACE2 were performed as described in Methods. As can be seen from Figure 5, the plant-produced RBD variants and S1 protein demonstrated specific binding to AEC2, the SARS-CoV-2 receptor. Notable, Endo deglycosylated RBD exhibited stronger binding to AEC2 compared with glycosylated counterpart.

**Figure 5.**
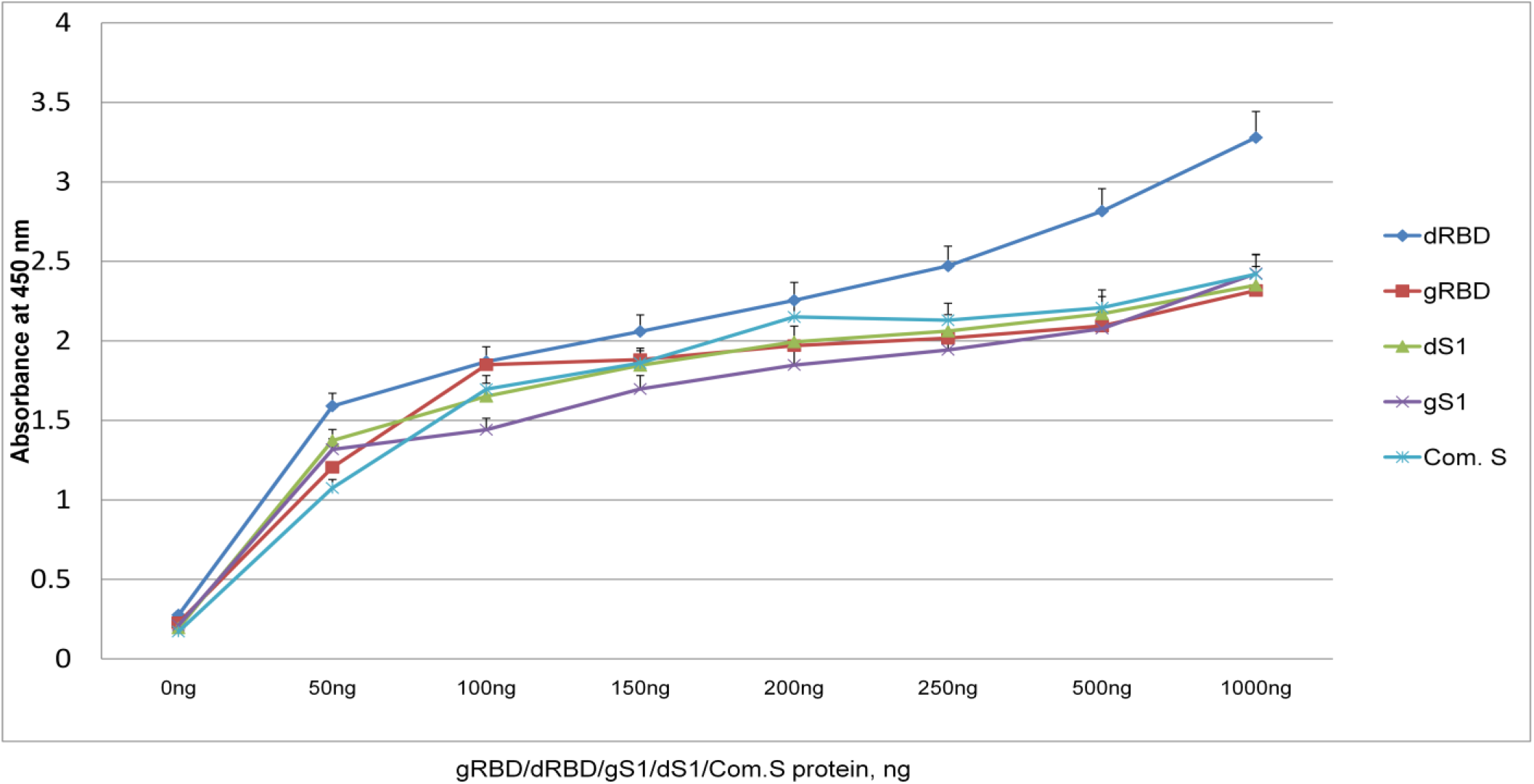
Binding affinity of gRBD, dRBD and gS1 to ACE2, receptor of SARS-CoV-2. Plant produced gRBD, dRBD, gS1 variants and baculovirus-Insect cells produced, commercially available recombinant spike protein of SARS-CoV-2 were incubated on plates coated with ACE2. After incubation, COVID 19 Spike RBD Polyclonal Antibody were added into each welland detected with rabbit IgG conjugated with HRP.

**Figure 6.**
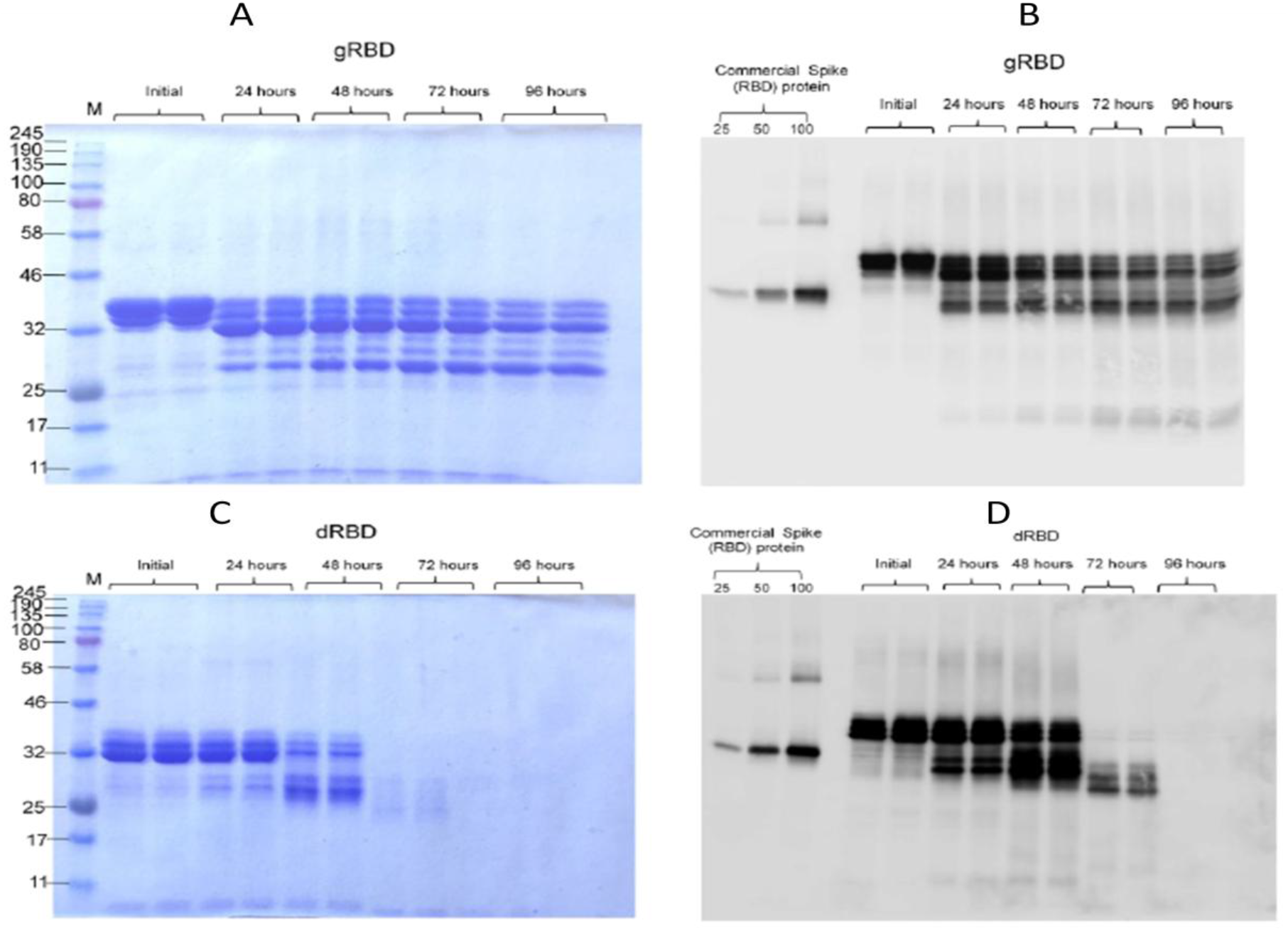
Stability of glycosylated and deglycosylated RBD variants. Plant produced, FLAG antibody affinity column purified glycosylated (gRBD, A, B) or *in vivo* Endo Hdeglycosylated (C, D) dRBD variants were incubated at 37 °C for 24, 48, 72 and96hours, and analyzed in SDS-PAGE (A, C) and Western blotting (B, D). Lanes were loaded with ~4.0 μggRBD (A) or dRBD (C). M: color prestained protein standard.Lanes were loaded with ~200 ng gRBD (B) or dRBD (D). Proteins on the blotwere probed with COVID 19 Spike RBD Polyclonal Antibody. The image was taken using the highly sensitive GeneGnomeXRQChemiluminescence imaging system.

**Figure 7.**
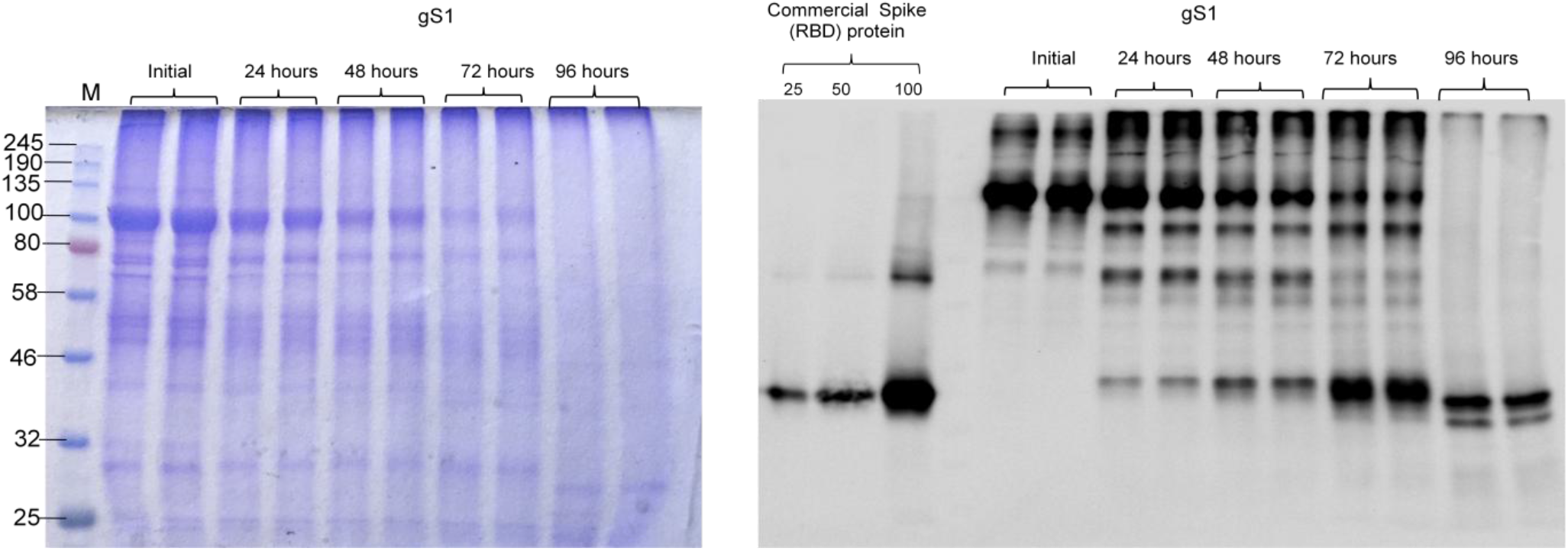
Stability of glycosylated S1 protein. Plant produced, Ni-NTA Resin affinity column purified glycosylated S1 protein was incubated at 37 °C for 24, 48, 72 and96 hours, and analyzed in SDS-PAGE (A) and Western blotting (B). Lanes were loaded with ~5.0 μg (A) or 200 ng (B) S1. M: color prestained protein standard. Proteins on the blot were probed with COVID 19 Spike RBD Polyclonal Antibody (B). The image was taken using the highly sensitive GeneGnome XRQ Chemiluminescence imaging system.

**Figure 8.**
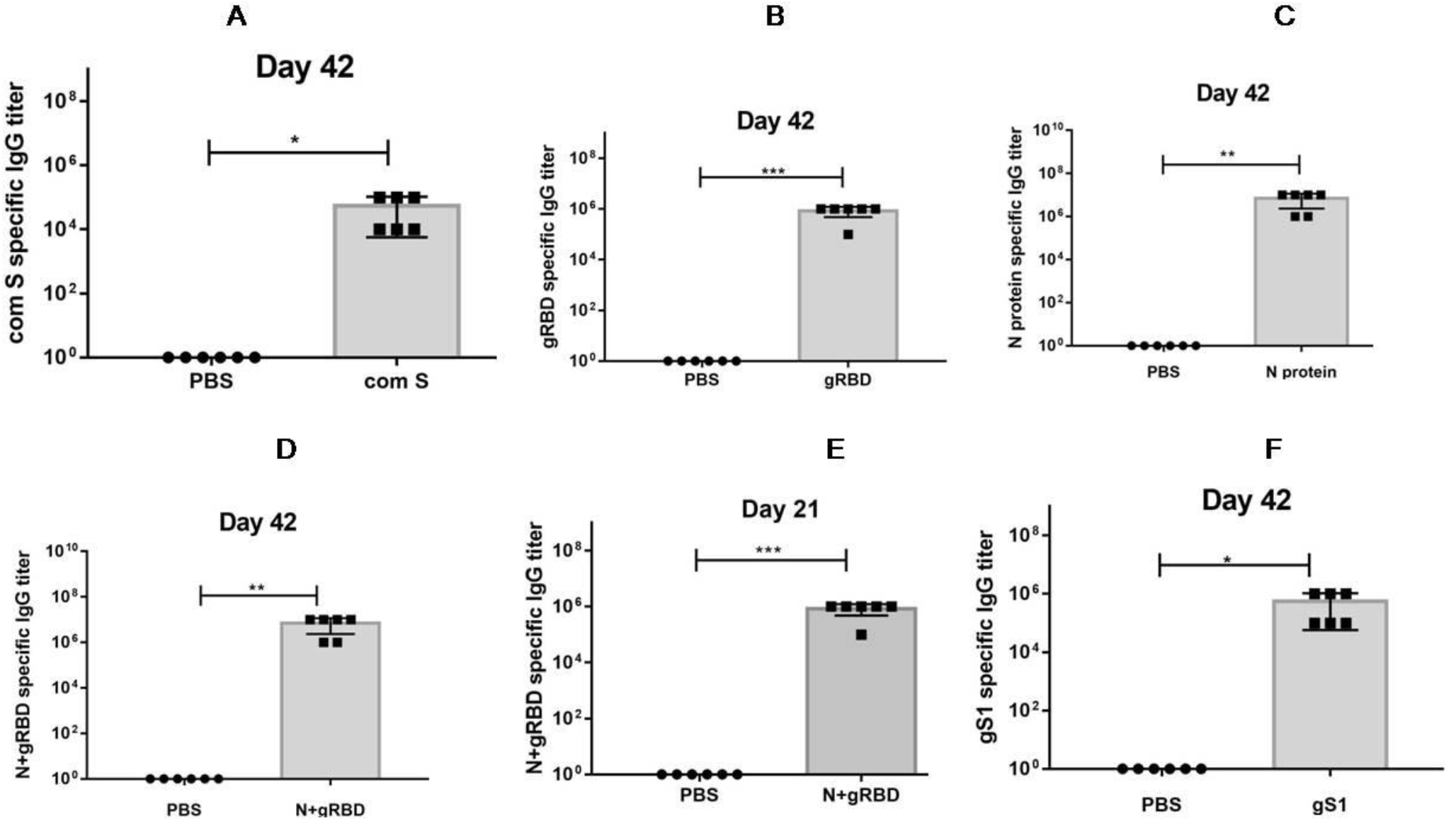
Immunogenicity of plant produced N and S variants in mice. Mice were immunized on study days 0 and 21 IM with 5 μg (with alum) of plant produced N and S variants. IgG responses in mice elicited by plant produced N, gRBD, N+gRBD and gS1 variants using Alhydrogel as an adjuvant. IgG titers were determined by ELISA in sera collected on 21st or 42nd days post vaccination from mice immunized with N and S variants. Detection of IgG titers in the mice serum with different dilutions specific to commercial available S protein (A), gRBD (B), N (C), N+gRBD (D, E) and gS1 (F) at day 21 (E) or 42 (A, B, C, D, F). Each point on the graph were derived from three replica for each dilution.*P < 0.05; **P < 0.01; ***P < 0.001(n=6 mice/group).

### Stability assessment of RBD variants and S1

The stability analysis of plant produced glycosylated and Endo H *in vivo* deglycosylated forms of RBD and gS1 of SARC-CoV were examined after incubation at 37 °C for 48, 96, 72 and 96 hours. The procedure of stability analysis was similar to those described previously^21^. These analyses showed that dRBD degradation was very low at incubation at 37 ° C for 24 hours (less than 10 %), however, at the same condition, gRBD degraded significantly (more than 60%). Plant produced gS1 protein showed better stability compared to gRBD or dRBD and degraded about 50% after 48 hours. Since the yield of Endo H dS1 was very low, we were not able to compare the stability of gS1 with dS1 counterpart.

### Immunogenicity studies of N, RBD and S1 variants in mice

Mice received two doses from each variant (N, gRBD, dRBD, [N+ RBD (cocktail)] or gS1 proteins) adsorbed to 0.3% Alhydrogel at three-week intervals (0, 21 days). Serum samples were collected on days −1 (pre-bleed) and 42 (post 2nd vaccination) and assessed for anti-N, anti-gRBD, anti-dRBD anti-(N+gRBD) and anti-gS1 antibody responses by an IgG ELISA. IgG responses showed that plant produced N and S protein variants were able to induce significantly higher titers of antibody with alum adjuvant at the 5 μg dose (Figure 6).

## Discussion

SARS-CoV-2 is a novel and highly pathogenic coronavirus, which has caused an outbreak in Wuhan city, China in 2019, and then spread rapidly throughout the world. The new coronavirus COVID-19 disease is currently responsible for the pandemic, created huge global human health crisis with significant negative impacts on health and economy worldwide. The development of effective and safe vaccines is urgently needed to prevent the spread of the disease and protect populations. A number of different types of vaccines (adenovirus-vectored vaccine, mRNA vaccines, inactivated vaccines and protein based subunits vaccines) are currently in various developmentstages^30–32^ and Phase 1 and 2 clinical trials have already been completed for some of them^5,33–36^. Several of these vaccine candidates have been demonstrated to induce antibodies specific for S and RBD and can neutralize SARS-CoV-2 infection^3^. However, due to725 mutations on SARS-CoV-2 S protein^37^, and considering possible future mutations on the S protein RBD, most of developed vaccines, especially mRNA and inactivated vaccines may not be effective against SARS-CoV-2. The other concern is if COVID-19 vaccines will be safe and effective. It was reported that SARS-CoV-2 RNAs can be reverse-transcribed and integrated into human genome^28^; this is one of the major concerns regarding the RNA based vaccines. The genomes of all known Coronaviruses including SARS-CoV, MERS-CoV and SARS-CoV-2 encode four structural proteins: N, S, E and M proteins. S protein plays a key role in the receptor (AEC2) recognition and responsible for attachment to host receptor. Nucleocapsid (N) protein, plays essential role in the viral genomic RNA packaging and facilitate the binding of the viral genome to a replication-transcription complex^39^. As mentioned above, most vaccine candidates such as DNA, mRNA, and viral vaccines developed so far are based on the S protein gene of SARS-CoV-2^4–7^. However, N Protein of SARS–CoV-2 could be a potential target for COVID-19 vaccine development since N protein is more conserved and stable, in fact few mutations was observed over time^2,10,12–14,40–42^. It has also been recently shown that the crystal structure of the SARS–CoV-2 nucleocapsid protein is very similar to coronavirus N proteins, which were previously described^43^. Cong et al. (2020) using a mouse hepatitis virus model showed that nucleocapsid protein contributes to forming helical ribonucleoproteins during the packaging of the RNA genome, and thereby regulating the viral RNA synthesis during replication and transcription. A number of studies have shown the critical roles of N protein for multiple steps of the viral life cycle^14^. It was also demonstrated that N proteins of many coronaviruses are highly immunogenic and are produced abundantly during infection^44^ and high levels of IgG antibodies against N protein have been detected in sera from patients who had recoveredfromSARS^45^. We performed sequence analysis and amino acid variations of structural proteins deduced from new coronavirus SARS-CoV-2 strains, isolated in different countries. Our amino acid sequence analysis of structural proteins demonstrates that, despite a high number of amino acid variations in membrane protein (M), there are no significant amino acid variations observed in the structural N, S or E-proteins obtained from new strains of SARS-CoV-2 coronavirus isolated in different countries^2^. These results support the importance of subunit vaccines that can be effective against SARS-CoV-2 worldwide, especially against various mutations. Notable, due to their safety, efficacy and scalability, subunit protein-based vaccines are becoming preferred alternatives to killed or live attenuated pathogens and are increasingly becoming popular. Plant transient expression system is a promising expression platform for production a variety of recombinant proteins such as vaccines, antibodies, therapeutic proteins and enzymes. Using a transient expression system, a number of difficult-to-express proteins have been successfully produced in *N. benthamiana* such as full length Pfs48/45 of *Plasmodium falcipatrium*^19,21^, human Factor IX and Furin, heptamerized form of PA63 of *Bacillus anthracis*^21^. Major challenge in any expression system, including plant expression systems, is to achieve the production of recombinant proteins with native folding, native like post-translational modifications (PTM), with high solubility and high yield. Flexible approaches are required for successful production of functional active recombinant proteins in plants with high yield. Thus, plant expression system could be ideal platform for rapid, cost effective, safe and high level production of structural proteins of SARC-COV-2. In this study, we demonstrate the production of functionally active structural proteins SARC-CoV-2, such as RBD variants, S1 domain and nucleocapsid proteins, at a high level and in a highly soluble form in the *N. benthamiana* plant using a transient expression platform. We purified these protein using anti-Flag and Ni-NTA resin chromatography. The purification yields were at least 20 mg/ kg plant leaf. In this study, we constructed two plasmids, expressing FLAG tagged and His tagged variants of RBD. We found that the expression level and purification yield of RBD-His tagged was low (less than 10 mg/kg leaf biomass), however, the expression level and purification yield of plant produced RBD-FLAG tagged protein was significantly higher, more than 20 mg/kg leaf biomass. The purified plant produced N and S proteins were recognized by N and S protein specific monoclonal antibodies demonstrating specific reactivity of mAb to plant produced N and S protein variants. The plant-produced RBD and S1 protein showed specific binding to AEC2, SARS-CoV-2 receptor. Notable, plant produced Endo H deglycosylated RBD exhibited stronger binding to AEC2 compared with glycosylated counterpart, suggesting native-like folding of dRBD, and negative effect of glycosylation on masking of important epitopes of the RBD.

IgG responses showed that plant produced N, S1 antigens were able to induce significantly higher titers of antibodies with Alhydrogel adjuvant. In this study, we are also reporting for the first time the co-expression of gRBD with protein N to produce a cocktail of SARS-CoV-2 antigens, which elicited high-titer antibodies compared to RBD or N proteins. Thus, N, S1 and [N+gRBD] antigens are promising vaccine candidates against COVID-19.These plants produced antigens can also be used as a diagnostic reagent in serological tests for detection of SARS-COV-2 antibody in COVID-19 patients.

## Methods

### Cloning and expression of Nucleocapsid(N) and Spike (S) protein variants in *N. benthamiana*

The nucleocapsid gene (1-419 AAGenBank accession YP_009724397) and Spike gene of SARS–CoV-2 variants: RBD (receptor-binding domain containing fragment, RBD, 319-591AA, GenBank accession MN985325, as a Flag or His6 tagged) and S1 domain (AA 14-815, GenBank accession MN985325, as a His6 tagged) were optimized for expression in *N. benthamiana* plants and de-novo synthesized at Biomatik Corp. To transiently express N, S protein (RBD, S1 domain and N+RBD) variants in *N. benthamiana*plants, the *Nicotiana tabacum* PR-1a signal peptide (MGFVLFSQLPSFLLVSTLLLFLVISHSCRA) was added to the N-terminus of N, RBD and S1 proteins. In addition, the KDEL sequence (the ER retention signal) and the FLAG epitope (the affinity purification tag for N and RBD proteins) or His6 tag (for S1 domain) were added to the C-terminus. The resulting sequences were inserted into the pEAQ^46^ binary expression vectors to obtain pEAQ-N, pEAQ-RBD and pEAQ-S1. These plasmids were then transferred into Agrobacterium AGL1 strain. To express N, RBD, S1 and N+RBD variants in *N. benthamiana* plant, AGL1 harboring pEAQ-N, pEAQ-RBD and pEAQ-S1 plasmids were infiltrated into *N. benthamiana* plant leaves. Plants were harvested at 4 dpi (day afterpost infiltration).

### Expression screening of N and S variants produced in *N. benthamiana* plant by Western blot analysis

SDS-PAGE analysis of plant produced N, RBD and S1 variants were performed on 10% acrylamide gels stained with Coomassie (Gel Code Blue, Pierce Rockford, IL). Western blot analysis was performed after electrophoresis and transfer of the proteins to Polyvinylidene Fluoride membranes. After transfer, Western blot membranes were blocked with I-Block (Applied Biosystems, Carlsbad, CA) and recombinant proteins N, RBD, S1 and N+RBD were detected with anti-DYKDDDDK antibody (cat. no. 651503, BioLegend, for N, RBD or N+RBD), anti-His antibody (for S1) and anti-RBD of S protein of SARC-CoV-2 monoclonal antibody (cat. no. MBS7135930, for RBD) or Human Novel Coronavirus Nucleoprotein (N) (1-419aa) monoclonal Antibody (MyBioSource, cat. no. MBS7135930).

### In vivo co-expressing of N and S protein with Endo H

To produce deglycosylated variants of N, RBD and S1 domain, AGL1 strain harboring pEAQ-N, pEAQ-RBD and pEAQ-S1 were co-infiltrated with pGeen-Endo H construct^19^.

### Purification of plant produced N, RBD and N+RBD (cocktail) proteins using anti-DYKDDDDK affinity gel

Purification of plant produced N and RBD (glycosylated and deglycosylated) variants were performed by anti-FLAG affinity chromatography using anti-DYKDDDDK affinity gel (cat. no. 651503, BioLegend) as described previously^21^. For purification, 20 g of frozen leaves, infiltrated with the pEAQ-N-Flag-KDEL or pEAQ-RBD-Flag-KDEL (with or w/o pGreen-Endo H) constructs were ground in 20 mL PBS buffer (1XPBS, 150 mMNaCI) using a mortar and a pestle. Plant debris was removed by filtration through Miracloth followed by centrifugation at20,000 g for 25 minutes and then filtered through a 0.45 μm syringe filter (Millipore). An anti-FLAG affinity column was prepared according to the manufacturer’s instructions. Sixty milliliters of a clear supernatant were loaded into 0.5 ml resin column equilibrated with PBS buffer. The column was washed with 10 volumes of PBS buffer. Bound proteins were eluted using 200 mM Glycine, 150 mM NaCl, pH 2.2 into tubes containing 2.0 M Tris solution to neutralize. Total protein content was estimated using the BioDrop and then analyzed by SDS-PAGE and western blot.

### Purification of plant produced S1 domain variant using HisPur™ Ni-NTA Resin

Plant produced S1 domain variants (glycoslated and deglycosylated variants) were purified using HisPur™ Ni-NTA Resin (ThermoFisher scientific, Cat. No. 8822) as described previously^19^. Briefly, 25 grams of frozen plant leaves, infiltrated with the pEAQ-S1-His6-KDEL construct, were ground using mortar and pestle in a phosphate extraction buffer (20 mM sodium phosphate, 300 mM sodium chloride, 10 mM imidazole, pH 7.4). The plant cell extract was filtered through Miracloth and centrifuged at 20,000 g for 25 minutes and then filtered through a 0.45 μm syringe filter (Millipore). The clarified total protein extract was then purified using HisPur Ni-NTA resin (Cat. No. 8822, Thermo Fisher Scientific) by following a previously used protocol^19^.

### Binding affinity of the plant produced RBD variants S1 to ACE2

Briefly a 96-well plate (Greiner Bio-One GmbH, Germany) was coated with 100 ng of commercially available ACE2 (MBS2563876, MyBiosource, USA) using 100 mM carbonate buffer and incubated overnight at 4°C. After incubation, the plates were blocked with blocking buffer for 2 h at room temperature. After blocking, various concentrations (50-1000 ng) of plant produced gRBD, dRBD, gS1 variants and baculovirus-Insect cells produced, commercially available recombinant Spike Protein of SARS-CoV-2 (RBD, His Tag, Arg319-Phe541, MM~ 25 kDa, MBS2563882, MyBioSource, USA) was added and incubated for 2 h at 37 °C. After 2 h, COVID 19 Spike RBD Coronavirus (COVID-19) Polyclonal Antibody was added into each well. The plate was washed three times with blocking solution (200 μl/well). After washing, wells were incubated with anti-rabbit IgG antibody (Cat. no. MBS440123, MyBioSource, USA). The plate was washed three times with 1X PBST washing solution (200 μl/wellfor 5 minute). 200 μl of Substrate Solution (Sigma) was added to each well. Afterwards plate was incubated in the dark, for 30 minutes at RT. After the incubation period, the plate was read at 450 nm on a multi well plate reader.

### Stability assessment of plant produced RBD variants and S1protein

The stability analysis of plant produced gRBD, dRBD and gS1 of SARS-CoV-2 recombinant proteins were examined after incubation at 37 °C for 48, 96, 72 and 96 hours using similar procedure as described previously^21^. Protein samples were diluted to 0.5 mg/ mL with PBS and were transferred into polypropylene Eppendorf low-binding tubes. After incubation at at 37 °C for 48, 96, 72 and 96 hours, samples were analyzed by SDS-PAGE and western blotting. The degradation of gRBD, dRBD or gS1 protein bands were calculated and quantified based on SDS-PAGE and WB analysis using highly sensitive Gene Tools software (Syngene Bioimaging, UK) and ImageJ software as described previously^21^.

### Immunogenicity studies of N, RBD, N+RBD and S1 domain in mice

Mice received two doses of N, RBD, N+RBD and S1 domain proteins adsorbed to 0.3% Alhydrogel at three-week intervals (0, 21 days). Groups of seven-week-old mice (6 animals/group) were immunized IM with 5 μg of N, RBD, N+RBD and S1 domainvariants at 0 and 21 days. IgG titers were determined by ELISA in sera collected on 21st or 42nd days post vaccination from mice immunized with N, S1 or N+ S1 (cocktail) variants.Serum samples were collected on days −1 (pre-bleed) and 42 (post 2nd vaccination) and assessed for anti-N, anti-RBD, anti-N+RBD and anti-S1 domain antibody responses by an IgG ELISA. Wells were coated with commercial Spike protein or plant produced RBD to detect antibody levels in serum of mice vaccinated with RBD or S1 protein. Wells were coated with plant produced N+ RBD cocktail protein to detect antibody levels in serum of mice vaccinated with cocktail N+ RBD protein.Wells were coated with plant produced N protein to detect antibody levels in serum of mice vaccinated with N protein.

Mice studies were carried out at Akdeniz University Experimental Animal CareUnit under permission of the Local Ethics Committee for Animal Experiments at Akdeniz University with thesupervision of a veterinarian.

GraphPad Prism software was used for all statistical analyses. Unpaired two-tailed t-test was used to compare antibody response of N, gRBD, gS1 and N+gRBD induced serums and PBS induced serums as control. Significant was accepted as P < 0.05 and P values were showed as *P < 0.05; **P < 0.01; ***P < 0.001. Each point on the graph were derived from three replica for each dilution.

### SDS-PAGE and Western blot analysis of purified N and S variants

SDS-PAGE analysis of plant produced N and S variants were performed on 10% acrylamide gels stained with Coomassie (Gel Code Blue, Pierce Rockford, IL). Western blot analysis was performed after electrophoresis and transfer of the proteins to Polyvinylidene Fluoride membranes. After transfer, Western blot membranes were blocked with I-Block (Applied Biosystems, Carlsbad, CA) and recombinant proteins detected with an anti-FLAG (N, S1 and N+S1) or anti-His Antibody (S2), or anti-SARS-COV2 COVID 19 Spike Protein Coronavirus Monoclonal Antibody (MyBioSource, cat. no. MBS2563837). The image was taken using highlysensitive GeneGnome XRQ Chemiluminescence imaging system (Syngene, A Division of Synoptics Ltd).

## Acknowledgments

The authors are grateful to Dr. George P. Lomonossoff (John Innes Centre, Biological Chemistry Department) and Plant Bioscience Limited for kindly providing pEAQ binary expression vector. This study was funded by the Akdeniz University to Tarlan Mamedov.

The authors are grateful to Dr. George P. Lomonossoff (John Innes Centre, Biological Chemistry Department) and Plant Bioscience Limited for kindly providing pEAQ binary expression vector. We thank Dr. Munevver Aksoy at Akdeniz University for editorial assistance.

## Author Contributions

T.M. Conceived the study; T.M. designed the experiments. D.Y., M.I., I.G., B.G, G.M., D.S. performed the experiments. T.M., G.H.. analyzed the data. T.M.., G.H. contributed to writing the paper.

